# Subcutaneous Neurotrophin-3 infusion induces corticospinal neuroplasticity and improvements in dexterity and walking in elderly rats after large cortical stroke

**DOI:** 10.1101/427864

**Authors:** Denise Duricki, Sotiris Kakanos, Barbara Haenzi, Christina Wayman, Nora Bödecker, Antje Vogelgesang, Diana Cash, Lawrence Moon

## Abstract

There is an urgent need for a therapy which reverses disability after stroke when initiated in a time frame suitable for the majority of new victims. Neurotrophin-3 (NT3) is a growth factor made by muscle spindles and skin which is required for the survival, development and function of locomotor circuits involving afferents from muscle and skin that mediate proprioception and tactile sensation. Its level declines in muscle and other tissues postnatally. We show that levels of NT-3 in the bloodstream were low in humans with ischemia stroke relative to young healthy controls. Accordingly, we set out to determine whether subcutaneous delivery of NT3 improves sensorimotor recovery after stroke in elderly rats. We show that one-month-long subcutaneous infusion of NT3 protein induces sensorimotor recovery after cortical stroke in elderly rats. Specifically, in a randomised, blinded pre-clinical trial, we show improved dexterity, walking and sensory function in rats following cortical ischemic stroke when treatment with NT3 is initiated 24 hours after stroke. Importantly, NT-3 was given in a clinically feasible time frame via this straightforward route. MRI and histology showed that recovery was not due to neuroprotection, as expected given the delayed treatment. Rather, anterograde tracing showed that corticospinal axons from the less-affected hemisphere sprouted in the spinal cord from cervical levels 2 to 8. Importantly, Phase I and II clinical trials by others show that repeated, subcutaneously administered high doses of recombinant NT-3 are safe and well tolerated in humans with other conditions. This paves the way for NT-3 as a therapy for stroke.

## Introduction

Stroke rapidly kills brain cells and is frequently disabling. Globally, there are 31 million stroke survivors, with another 9 million new strokes annually (WHO). The majority of stroke victims are not admitted to hospital and diagnosed within six hours ^1^ yet clot-busting therapies only work when treatment is initiated well within 4.5 hours. New therapies are urgently needed ^2,3^.

We are the first to study whether recombinant human neurotrophin-3 (NT-3) can improve recovery when given in a clinically-feasible time-frame after stroke. Others have shown that NT-3 plays a role in the development, function and repair of locomotor circuits ^4–8^ including reports that intracranial delivery of NT-3 immediately following stroke or by intracranial gene therapy prior to stroke reduces infarct volume ^9–11^. Moreover, NT-3 restores sensorimotor function following spinal cord injury in rats ^6,12–14^ by promoting axon growth and synaptic plasticity in multiple locomotor pathways including the corticospinal tract and proprioceptive pathways. All these systems express trkC and/or p75 receptors for NT-3 ^15–17^. We therefore examined the ability of NT-3 to promote recovery in a model of stroke.

We chose to deliver NT-3 by the subcutaneous route for translational relevance. First, high, repeated subcutaneous doses of recombinant NT-3 are safe and well-tolerated in Phase I and II clinical trials for other disorders ^18–21^. This is a clinically-straightforward route after ischemic stroke in humans. Peripherally administered neurotrophins accumulate in sensory ganglia where they can cause gene transcription, axon growth of primary afferents, and synapse strengthening within locomotor circuits ^8,22–27^. Because >90% of strokes occur in people older than 65 with comorbidities including carotid arterial stenosis^28^, we evaluated the effectiveness of NT-3 in elderly rats with large cortical strokes caused by distal middle cerebral artery occlusion and common carotid occlusion^29^. We chose to deliver our therapy 24 hours after cortical ischemia, because most stroke victims are admitted and diagnosed by this time^1,30^.

We show that delayed NT-3 improves sensory and motor recovery after large cortical stroke in elderly rats.

## Results

### Neurotrophin-3 levels in blood serum are low after stroke in elderly humans

It has been shown previously that Neurotrophin-3 levels decline generally in most tissues in the postnatal period ^28,29^. We now show that whereas the level of Neurotrophin-3 in serum was modest in young healthy adults, the level was low in people after ischemic stroke (Figure 1). Given these low levels, in the present study we sought to elevate the levels of Neurotrophin-3 after large cortical stroke in elderly rats.

**Figure 1:**
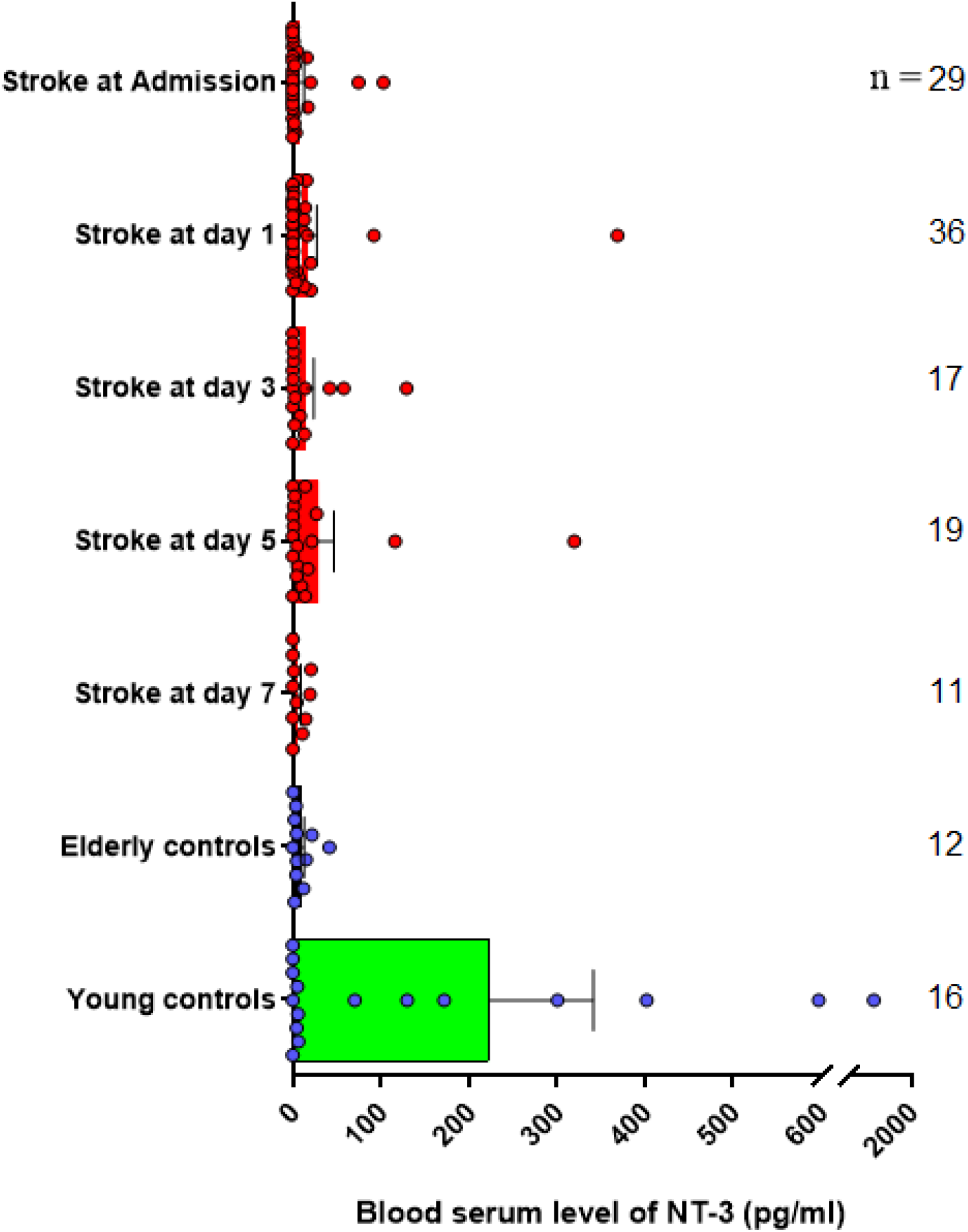
ELISA of serum shows that NT-3 levels were very low in humans on admission to hospital after ischemic stroke relative to young healthy controls (Kruskal-Wallis p=0.0543; Dunn’s p=0.033). The level of NT-3 was not increased up to seven days after ischemic stroke (versus stroke at admission; Dunn’s p-values >0.05). Numbers of subjects per group are shown on the graph. Other information can be found in Table 1. Graph shows means and SEMs.

Ischemic stroke was induced in the sensorimotor cortex representing each rat’s dominant forearm by permanent distal occlusion of the middle cerebral artery with tandem occlusion of the common carotid arteries ^29^ (Figure 2). Twenty-four hours later, rats were randomised to treatment with treatment allocations concealed using coded vials. Magnetic resonance imaging (MRI) was performed and then osmotic minipumps were implanted subcutaneously. Recovery of sensory and locomotor performance was assessed weekly for 12 weeks. Anterograde tracer was injected into the less-affected sensorimotor cortex 4 weeks before the end of each study. All surgeries and treatments were performed using a randomised block design and the experimenter was fully blinded to treatment allocation until all analyses were completed. The rats were euthanized at the end for histology.

**Table 1:** Patients and Controls. NIHSS: National Institute of Health Stroke Scale; * data available for 45 patients.

|  | age<br>(years)<br>range | age<br>(years)<br>median | sex m/f | NIHSS median | n |
| --- | --- | --- | --- | --- | --- |
| **Young controls** | 20-34 | 22 | 6/10 | n/a | 16 |
| **Elderly controls** | 70-87 | 81 | 6/6 | n/a | 12 |
| **Stroke patients** | 48-92 | 74 | 31/22 | 15* | 53 |

**Figure 2:**
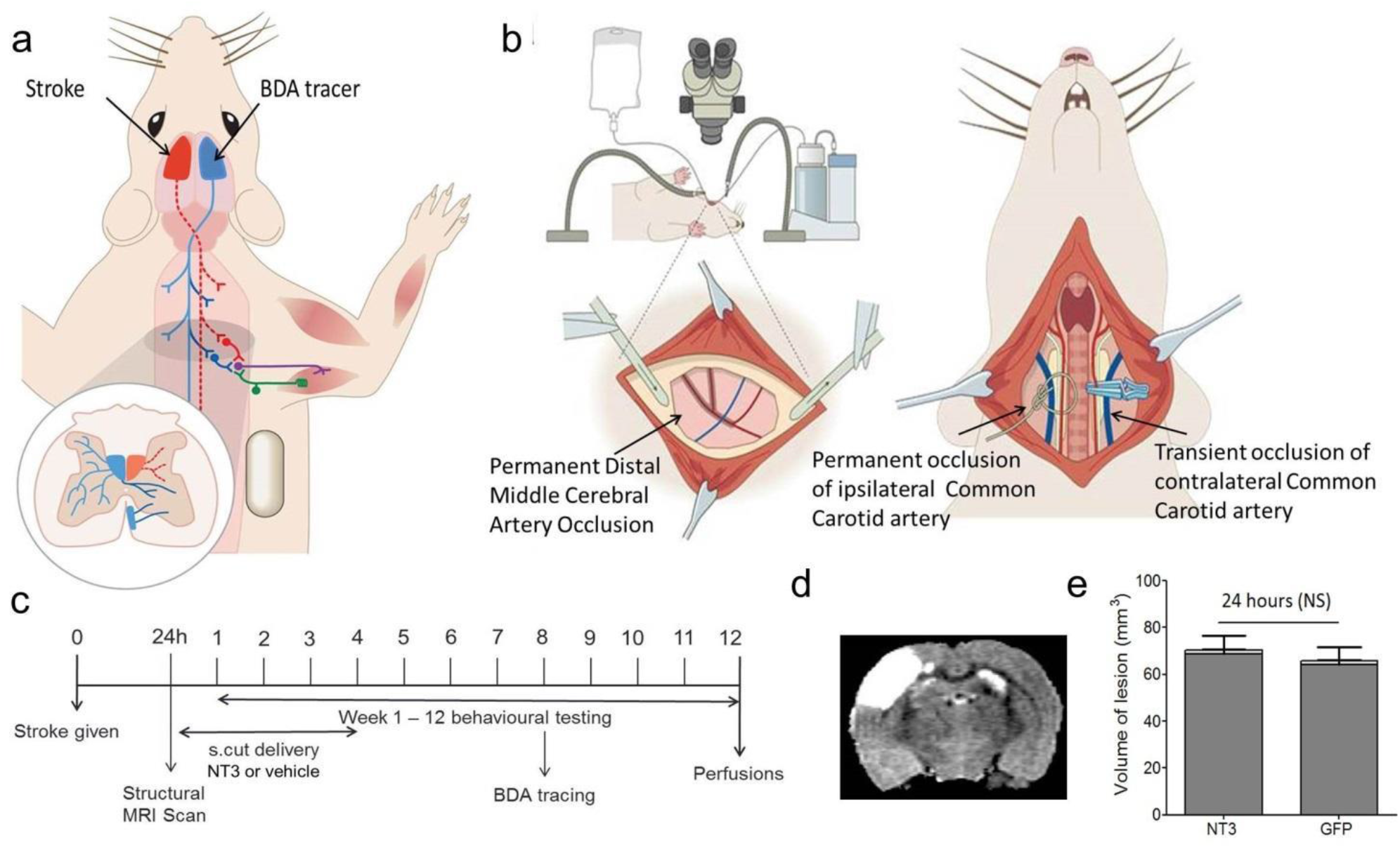
Delayed subcutaneous infusion of NT-3 did not reduce infarct volume in elderly rats. (A) Rats were randomised to treatment with recombinant NT-3 or vehicle from subcutaneous minipumps, implanted 24 hours after stroke (red). Behavioural testing was conducted for twelve weeks. The spared corticospinal tract was traced anterogradely (blue) eight weeks after onset of treatment. (B) MR images taken 24 hours after stroke show large cortical infarcts with no differences between groups (p<0.05).

### Delayed subcutaneous NT-3 did not neuroprotect

MRI at 24 hours after stroke (immediately before treatment) showed that large infarcts involved motor cortex and somatosensory cortex as well as cortex rostral and caudal to these regions (Figure 2). There were no differences between stroke groups in lesion volumes. Therefore, as expected from our previous work, stroke causes a focal infarct in sensorimotor cortex and loss of cortical efferents to the cervical spinal cord, and delayed treatment with NT3 does not cause neuroprotection (consistent with the delayed time frame of administration).

### Delayed subcutaneous NT-3 promoted sensory and motor recovery

We assessed recovery of dexterity using the staircase test. Prior to stroke, elderly rats retrieved more than 80% of pellets with their dominant paw and more than 60% of pellets with their other paw (Figure 3A). Stroke was induced in the hemisphere representing the dominant paw. One week after stroke, the two groups were impaired in reaching for pellets with their affected paw. Recovery was greater in the group of rats treated with NT-3 than those treated with vehicle (Figure 3A).

**Figure 3:**
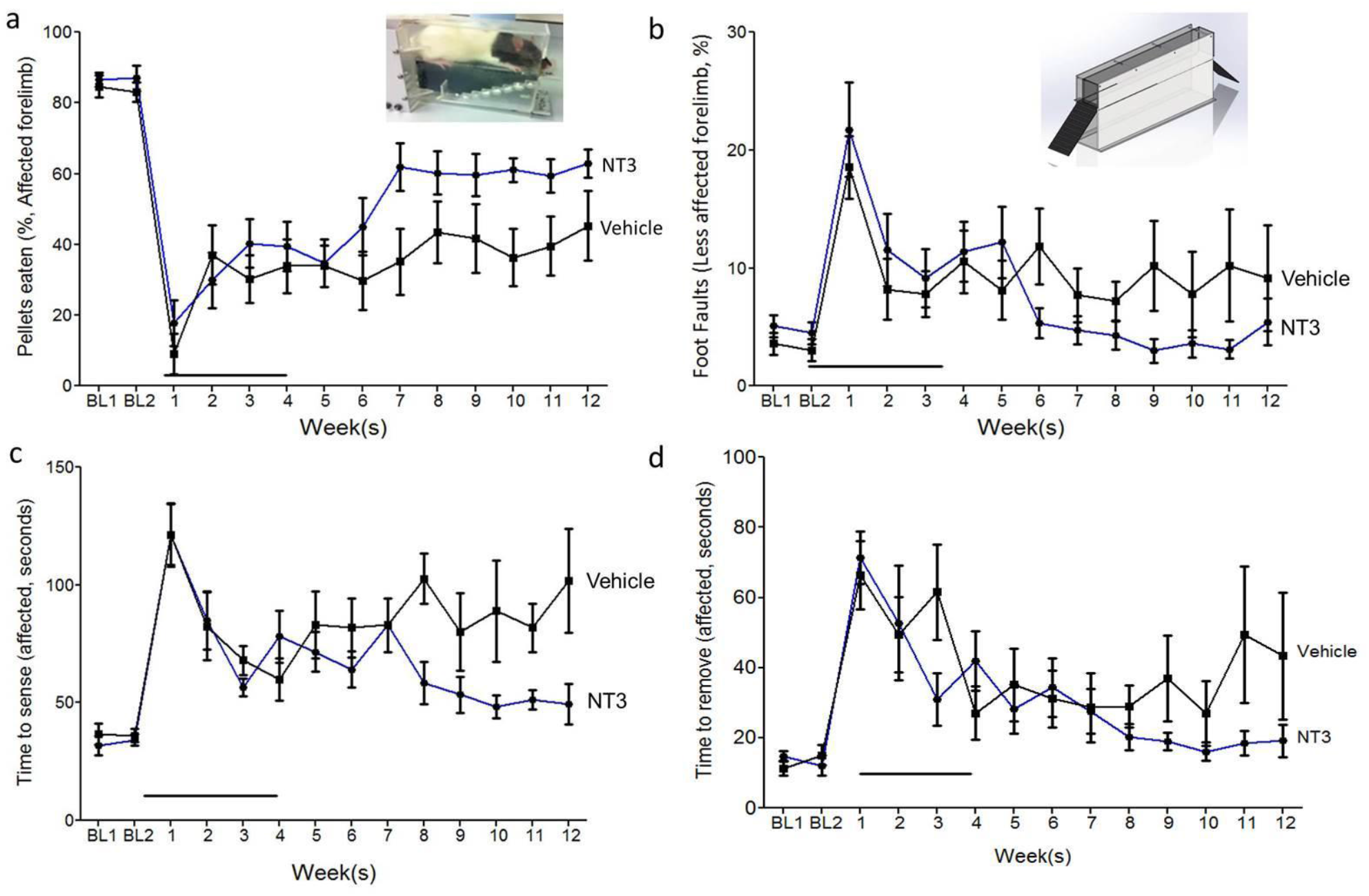
Delayed subcutaneous infusion of NT-3 improved dexterity. (A) The “staircase test” was used to assess dexterity. After stroke, the two groups of rats were similarly impaired in their ability to reach for pellets with their affected forelimb (t-test, p=0.18). Recovery was greatest in the group of rats treated with NT-3 compared to vehicle (treatment × time F_11,17_=2.91, p=0.024). (B) The “horizontal ladder” test was used to assess accuracy of foot placement. One week after stroke, the two groups of stroke rats made a similar, large number of errors with their affected forelimb (t-test, p=0.96). Recovery was nearly complete in rats treated with NT-3 compared to vehicle (treatment × time F_11,215_=2.33, p=0.01). (C) The “sticky patch test” was used to assess the time taken first to contact and then to (D) remove stimuli on their forelimbs. After stroke, the two groups of rats were similarly slow to contact a sticky patch stuck to their affected wrist (t-test, p=0.91). Recovery was greatest in the group of rats treated with NT-3 compared to vehicle (treatment F_1,21_=7.2, p=0.014; treatment × time F_11,20_=2.6, p=0.31).

We also assessed sensory responsiveness and dexterity using the sticky patch test. Stickers of equal radius were attached to each wrist and the time taken to contact and then remove them was measured (Figure 3B). Prior to stroke, rats contacted the first sticker rapidly and then removed it rapidly. One week after stroke, both groups took longer to contact the first sticker (Figure 3B). The stroke group treated with NT-3 recovered more fully than the stroke group treated with vehicle. Similarly, one week after stroke, both groups took longer to remove the first sticker (Figure 3C). The stroke group treated with NT-3 recovered more fully than the stroke group treated with vehicle, removing it more rapidly.

Finally, we assessed accuracy of limb placement during walking using a horizontal ladder with irregularly spaced rungs (Figure 3D). Accuracy of limb placement requires proprioceptive sensory feedback from the limbs especially when rung spacing was changed frequently to avoid fixed stride lengths or other pattern learning ^31^. Video recordings were assessed in slow motion to count the number of errors made with the affected forelimb (expressed as a percentage of steps). Prior to stroke, rats crossed the ladder making very few errors (Figure 3D). One week after stroke, both groups of rats made a large, comparable number of errors with their affected forelimb. The stroke group treated with NT-3 recovered more fully then the stroke group treated with vehicle (Figure 3D). In summary, delayed, subcutaneous treatment with NT-3 improved sensory and motor recovery after large cortical stroke.

### NT-3 induced neuroplasticity in the corticospinal tract

Anterograde tracer (biotinylated dextran amine, BDA) was injected into the less-affected sensorimotor cortex 4 weeks prior to the end of the study (Figure 2). Immunolabelling for BDA was used to assess axonal sprouting of this corticospinal tract. This decussates in the medullary pyramids and largely projects to the contralateral side of the spinal cord in the dorsal columns and in the dorsolateral columns; there is also a minor projection which does not decussate but runs in the ventral spinal cord (see inset in Figure 4). We chose to measure corticospinal tract axons in every cervical segment from C2 to C8 because the shoulder, forelimb and paw muscles are supplied by dorsal root ganglia (DRG) and motor neurons between C2 and C8 ^32,33^. Corticospinal axons from the less-affected hemisphere were measured at three parasagittal planes on the denervated side (“Midline”, “Distance 1” and “Distance 2”; Figure 3) and obliquely where ventral corticospinal axons that have projected ipsilaterally from cortex enter the grey matter (“Ipsi”; Figure 3). Figure 3 shows representative pictures of corticospinal axons at C7. Neurotrophin-3 treatment increased corticospinal tract sprouting at all levels from C2 to C8 (Figure 4). Sprouting was frequently observed at the midline, at distance 1, distance 2 and from the ventral corticospinal tract. Increased sprouting at distance 2 indicates that more corticospinal tract axons extended into the territory of the forelimb motor neuron pools which are located in lateral and ventrolateral grey matter (Figure 4) ^28,29^.

**Figure 4:**
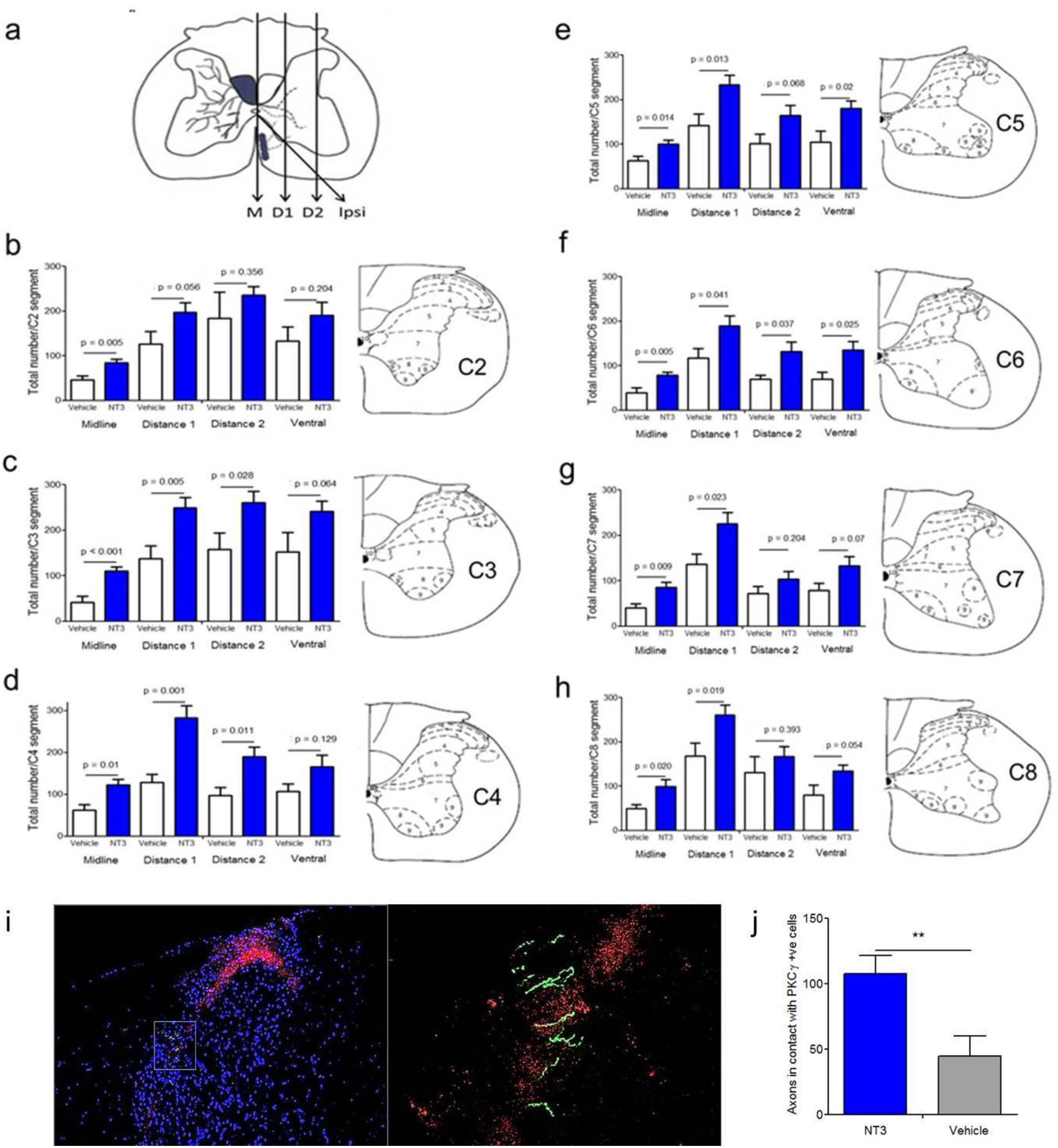
NT-3 treatment caused plasticity of corticospinal axons in the cervical spinal cord. A-H) Anterogradely traced corticospinal axons were counted in the cervical cord segments C2 to C8 at the midline (M), at two more lateral planes (D1 and D2) and crossing into grey matter from the ipsilateral, ventral tract (Ipsi). I) Corticospinal tract axons (green) sprouted into the dorsal horn on the affected side amongst PKC gamma labelled sensory neurons (red). J) NT3 enhanced the number of corticospinal tract axons which sprouted into the laminae containing PKC gamma labelled sensory neurons.

### NT-3 was transiently elevated in the bloodstream

ELISA showed that NT-3 was elevated in the serum of elderly rats treated subcutaneously with NT3 relative to those treated with vehicle (Figure 5) at 2 weeks after onset of treatment but not at 12 weeks (*i.e.*, 8 weeks after removal of minipumps). This was expected and is consistent with NT-3’s short half-life in serum^34^.

**Figure 5:**
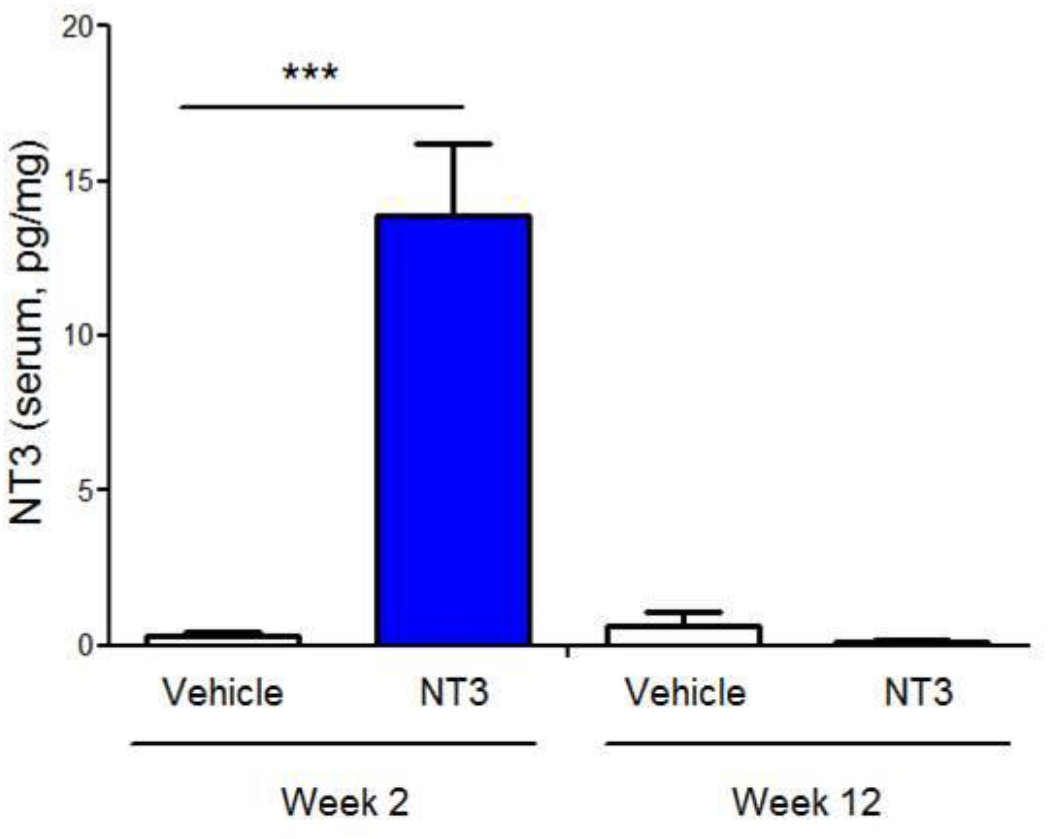
Subcutaneous infusion of NT-3 enhanced levels of NT-3 in serum of elderly rats. ELISA showed that serum levels of NT-3 were elevated at 2 weeks (unpaired t-test, p<0.001) but not 12 weeks (unpaired t-test, p>0.05).

## Discussion

We report that the level of Neurotrophin-3 in serum was modest in young healthy adult humans, whereas the level was low in elderly people and it remained low in elderly people after stroke whether measured at hospital admission or up to seven days later. A smaller study reported a very slight elevation in serum levels of NT-3 in patients with cortical infarcts ^33^. Together these studies provide a motivation for exploring whether supplementation of NT-3 improves recovery after stroke.

Subcutaneous treatment with NT-3, initiated 24 hours after stroke, caused changes in multiple spinal locomotor circuits, and promoted a progressive recovery of sensory and motor function in elderly rats. The fact that NT-3 can reverse disability even when initiated 24 hours after stroke is exciting because the vast majority of stroke victims are diagnosed within this time frame ^1^ (which is not the case for the gold-standard drug for stroke, tPA, which needs to be given within a few hours and is only available to a minority). Thus, NT-3 could potentially be used to treat an enormous number of victims.

NT-3 has good clinical potential. Firstly, it would be feasible to give hospitalised stroke patients subcutaneous infusions of recombinant NT-3. Secondly, Phase II clinical trials show that subcutaneous doses of NT-3 up to 150 μg/kg/day are well tolerated and safe in healthy humans and in humans with other conditions^18,19,21,35,36^, which paves the way for NT-3 as a therapy for stroke. We used a similar dose in this study: in future experiments, one might optimize the dose and duration of treatment because it is possible that a different dose of NT-3 would promote additional recovery after stroke, especially if combined with appropriate rehabilitation^37^. Secondly, there is good conservation from rodents to primates including humans in the expression of receptors for NT-3 in the locomotor system ^15,38–41^. Thirdly, in none of our rodent experiments has NT-3 treatment caused any detectable pain, spasticity or muscle weakness (in line with the human trials).

Our work is relevant to recent work shedding light on the mechanisms whereby sensory cues enhance recovery after CNS injury in humans and other animals^31,42^. It has been proven that proprioceptive signals cause sprouting of descending locomotor pathways and enhance recovery after unilateral spinal cord injury because genetic depletion of muscle spindles attenuates central plasticity and impairs recovery^31^. We now extend these findings by proving that NT-3 can cause sprouting of descending locomotor pathways and enhance locomotor recovery after unilateral stroke, even when onset of treatment is delayed by 24 hours. Thus, at least to some extent, subcutaneously infused NT-3 may mimic proprioceptive signalling after neurological injury. This idea is also consistent with work from postnatal mice showing that a single injection of NT-3 can restore monosynaptic connectivity between proprioceptive afferents and spinal motor neurons with genetically depleted muscle spindles ^23^. We have now begun a program of work designed to dissect the mechanisms whereby NT-3 promotes recovery. We are currently testing the hypothesis that NT3 enters the DRG and causes changes in gene expression in proprioceptive neurons in adult rats after stroke, as has been shown in development, which then drives central changes ^22^. We and others have already shown that neurotrophins accumulate in sensory or sympathetic ganglia after subcutaneous, intravenous, intramuscular or intraneural administration following transport in the bloodstream or retrograde transport in nerves ^27,43–46^. Many groups have shown that the dorsal root ganglia is highly vascularised and contain fenestrated endothelial cells which make its blood-nerve interface leaky to proteins as big as (or bigger than) neurotrophin-3. We now plan to use RNAseq to identify transcriptional changes in the DRG that might cause the functional recovery that we observe.

In summary, treatment of elderly rats with NT-3 (initiated in a clinically-feasible time-frame) induces multilevel corticospinal neuroplasticity, improves sensory function, walking and dexterity.

## Methods

Full Materials and Methods are available in Supplementary Information.

### Patients and Controls

Stroke Patients were recruited at the Stroke Unit of the University Medicine Greifswald, Germany. Venous blood was sampled upon admission to the emergency room within 12h of stroke onset and on the morning of the days 1, 3, 5 and 7 thereafter. Patients with age >18 years suffering from acute ischemic stroke were eligible for this study within 12 h of disease onset if their National Institute of Health Stroke Scale Score (NIHSS) was ≥ 6 and no signs of infection were detected (C-reactive protein ≤ 50 mg/L and procalcitonin ≤ 0.5 ng/mL). Patients were treated with standard medical care in a dedicated stroke unit and did not receive immune-suppressive drugs. Lysis by recombinant tissue plasminogen activator and thrombectomy were carried out as clinically indicated. Age-matched control individuals were either healthy or were recruited from the ophthalmology clinics from patients scheduled to receive cataract surgery. Young controls were healthy individuals. See Patient and control characteristics in Table 1.

### Ethics approval and consent to participate

The study protocol was approved by the ethics committee of the Medical Faculty, University of Greifswald (No. III UV 30/01). All donors provided written informed consent directly or through a surrogate.

### ELISA

To gain serum, 5ml of venous blood were taken into BD vacutainer serum sampling tubes (SST II advanced, Ref 367955, Beckton Dickenson, Heidelberg, Germany). Tubes were centrifuged at 3000g for 5 min at room temperature. Aliquots were stored at −80°C. NT-3 was measured by standard ELISA according to manufacturer’s instructions (ab100615 – Neurotrophin 3 human ELISA Kit, abcam, Cambridge, United Kingdom).

### Statistical analysis

Human serum data sets were tested for adherence to the Gaussian distribution with the Kolmogorov-Smirnov test. Since the data failed the normality test, we used non-parametric testing throughout. The Kruskal-Wallis test with Dunn’s multiple comparison test as a post- test were used as appropriate. GraphPad-PRISM 5.0 (GraphPad Software inc., San Diego, CA, USA) was used for all analyses. A p-value < 0.05 was regarded as significant.

#### Animal experiments

53 elderly rats (aged ~20 months; Charles River, UK) were trained on behavioural tasks for 4 weeks prior to baseline scoring. Stroke was induced unilaterally in the sensorimotor cortex by permanent distal middle cerebral artery occlusion (MCAO) and tandem carotid occlusion, as described elsewhere ^29^. Twenty-four hours later, recombinant NT-3 or vehicle was infused subcutaneously for four weeks. Anaesthesia and analgesia were used. Rats were assessed for 12 weeks using the staircase test, horizontal ladder, and sticky patch tests. All surgeries, treatments and behavioural testing were performed blind using a randomized block design. Group codes were only broken after all analyses were completed. n=9-12/group were used for tract tracing. Behavioural data were analysed using linear models using baseline scores as covariates. Anterograde tracing data were assessed using one way ANOVA and *post hoc* t-tests. Threshold for significance was 0.05. Two-tailed tests were used. Data are presented as means ± SEM. Asterisks indicate *p ≤ 0.05; **p ≤ 0.01; *** p ≤ 0.001.

## Acknowledgments

The research leading to these results has received funding from the European Research Council under the European Union’s Seventh Framework Programme (FP/2007-2013) / ERC Grant Agreement n. 309731 and by a “Serendipity Grant” from The Dunhill Medical Trust (grant number: SA21/0512). The NT3 ELISA was funded by Brain Research UK (201617-04) and the Medical Research Council (S026053/1 to LDFM). AV received funding from the University of Greifswald (Käthe-Kluth-Research Group). The funders had no role in the study design, data collection and the analysis, decision to publish, or preparation of the article. The funders had no role in the study design, data collection and the analysis, decision to publish, or preparation of the paper.

